# Chronic methamphetamine administration produces cognitive deficits through augmentation of GABAergic synaptic transmission in the prefrontal cortex

**DOI:** 10.1101/2022.02.28.482382

**Authors:** Monserrat Armenta-Resendiz, Ahlem Assali, Evgeny Tsvetkov, Christopher W. Cowan, Antonieta Lavin

## Abstract

**BACKGROUND:** Chronic methamphetamine (METH) abuse is associated with the emergence of cognitive deficits and hypofrontality, a pathophysiological marker of many neuropsychiatric disorders that is produced by altered balance of local excitatory and inhibitory synaptic transmission. However, there is a dearth of information regarding the cellular and synaptic mechanisms underlying METH-induced cognitive deficits and associated hypofrontal states.

**METHODS:** Rats went through a METH sensitization regime or saline (SAL) consisting of 14 days of METH treatment (day 1 and 14, 1 mg/kg; days 2-13, 5 mg /kg) followed by 7-10 days of home cage abstinence. Temporal Order Memory and Working Memory tests, chemogenetic experiments as well as whole-cell patch recordings on prelimbic PFC ex vivo slices were performed during abstinence.

**RESULTS:** We find here that repeated METH administration in rats produces deficits in working memory and increases in inhibitory synaptic transmission onto pyramidal neurons in the prefrontal cortex (PFC). The increased PFC inhibition is detected by an increase in spontaneous and evoked inhibitory postsynaptic synaptic currents (IPSCs), an increase in GABAergic presynaptic function, and a shift in the excitatory-inhibitory balance onto PFC deep-layer pyramidal neurons. We find that pharmacological blockade of D1 dopamine receptor function reduces the METH-induced augmentation of IPSCs, suggesting a critical role for D1 dopamine signaling in METH-induced hypofrontality. In addition, chronic METH administration increases the intrinsic excitability of parvalbumin-positive interneurons, a key local interneuron population in PFC that controls inhibitory tone. Using a cell type-specific chemogenetic approach, we show that increasing PV+FSI activity in the PFC is necessary and sufficient to cause deficits in temporal order memory similar to those induced by METH.

**CONCLUSION:** Together, our findings reveal that chronic METH exposure increases PFC inhibitory tone through a D1 dopamine signaling-dependent potentiation of inhibitory synaptic transmission, and that reduction of PV+FSI activity can rescue METH-induced cognitive deficits, suggesting a potential therapeutic approach to treating cognitive symptoms in patients suffering from methamphetamine use disorder.

## Introduction

Methamphetamine (METH) use disorder is a profound medical and societal problem. METH users frequently develop hypofrontality, a state of reduced frontal cortex activity associated with deficits in temporal order memory, working memory (WM), attention, and impulsivity (1–3). METH-induced cognitive deficits can significantly reduce the quality of life and contribute to sustained drug-taking and relapse (4,5). In addition, METH-induced cognitive impairments are long-lasting, with chronic METH users reporting cognitive dysfunction even after 11 months of abstinence (6,7). However, the mechanisms underlying METH-induced hypofrontality and cognitive dysfunction during abstinence remain elusive. Along with classic substance use disorder-like behaviors, such as drug craving and seeking, rodents repeatedly exposed to METH also exhibit cognitive abnormalities during abstinence (8–10), including memory and WM deficits (9–11). As such, repeated psychostimulant administration in rodents represents a strong, face-valid model to study the mechanisms that underlie hypofrontality and cognitive disabilities in METH use disorder.

Although inhibitory interneurons constitute only 10-20% of the neuronal population in the cortex(12), their function is critical to control the activity level of the principal glutamatergic pyramidal neurons (PNs) and to maintain the proper balance between excitatory (E) and inhibitory (I) synaptic transmission. E-I ratio alterations in the frontal cortex are thought to contribute to diverse neuropsychiatric disorders (13,14), including substance use disorders(15–19). The role of cortical glutamatergic excitatory neurons in cognition has been extensively studied (20–28), but we are only beginning to understand the contribution of cortical interneurons to cognitive abilities. Among the multiple GABAergic interneuron subtypes within the medial prefrontal cortex (mPFC) (12,29–31), the parvalbumin-positive fast-spiking interneurons (PV+FSIs) exert a particularly powerful influence on PN inhibitory tone. Indeed, PV+FSIs are the most common interneuron subtype(32), they contact nearly every local PN (33), and they target PNs near the site of action potential initiation (12,29). PV+FSI dysfunction has been linked to numerous neuropsychiatric disorders, such as epilepsy, schizophrenia, depression, autism, and Alzheimer’s disease (34–38). Importantly, PV+FSIs are particularly affected in patients with WM deficits(39–41). PV+FSIs have elicited interest for their critical role in orchestrating cortical oscillations, maintaining E-I balance, and mediating a variety of cognitive behaviors, including WM (41–44). However, while it is well known that repeated METH exposure induces cognitive dysfunction in both rodents and humans(6,7,9–11), its synaptic and cellular effects on PV+FSIs, and the mechanisms underlying METH-induced cognitive deficits, remain elusive. Since METH is known to increase dopamine (DA) release in the brain, including in the mPFC (45–47), DA receptor signaling could, at least in part, regulate changes in PFC activity, and consequently elicit PFC-dependent behavioral deficits.

VTA neurons project to both PFC PNs and PV+FSIs in layers I–III and V–VI (48–50), but several studies have shown that DA in the PFC preferentially acts on PV+FSIs (50–52). In PFC layers V-VI, PV+FSIs express high levels of dopamine D1 receptors (D1R) and low levels of dopamine D2 receptors (D2R) (31). Activation of D1R signaling on PV+FSIs increases their excitability – enhancing both evoked and spontaneous inhibitory post-synaptic currents (IPSCs) recorded in PNs (51,53–56).

Multiple studies demonstrate that DA release and GABA interneurons influence WM via their contribution in tuning recurrent excitation in mPFC networks (53,57–61). In addition, WM delay cells, which are mainly PN neurons that show spiking during memory delays of WM tasks (62–64), are modulated by DA activation of D1R (but not D2R) signaling, and optimal levels of D1R stimulation are essential for WM function (53,57,60,61).

In this study, we show that repeated METH administration produces hypofrontal-like states and cognitive deficits that are the consequence of a maladaptive, D1R-dependent enhancement of GABAergic transmission. Moreover, using cell type-specific chemogenetics, we demonstrate that reduction of PV+FSI activity is sufficient to rescue the cognitive deficits produced by chronic METH administration, suggesting a circuit-specific strategy to treat cognitive symptoms in individuals suffering from METH use disorder.

## Materials and Methods

### Animals

Long Evans male rats expressing Cre in parvalbumin-positive cells (PV+), obtained from NIDA (LE-Tg(Pvalb-iCre)2Ottc), were bred and maintained on a Long Evans background. Rats were housed under 12 light/12 dark hour cycle in pairs, with access to food and water ad libitum. All experiments were performed in accordance with the National Institutes of Health guidelines for the care and use of laboratory animals. Procedures were approved by the Institutional Animal Care and Use Committee (IACUC) of the Medical University of South Carolina.

### METH administration

Day 1 and day 14 of the treatment consisted of a single i.p injection (0.1ml/100g) of METH at 1 mg/kg or saline (SAL). From day 2 to day 13, rats received i.p injections (0.1ml/100g) of METH at 5 mg/kg or SAL. Following METH or SAL treatment, rats were given 7-10 days of home cage abstinence. One group of animals was tested for Temporal Order Memory (TOM), followed by electrophysiological recordings (Fig 1A), and a second group of animals was tested for working memory using a Delay Non-Match to Sample Task (DNMST; Fig. 1A), also followed by electrophysiological recordings.

**Figure 1.**
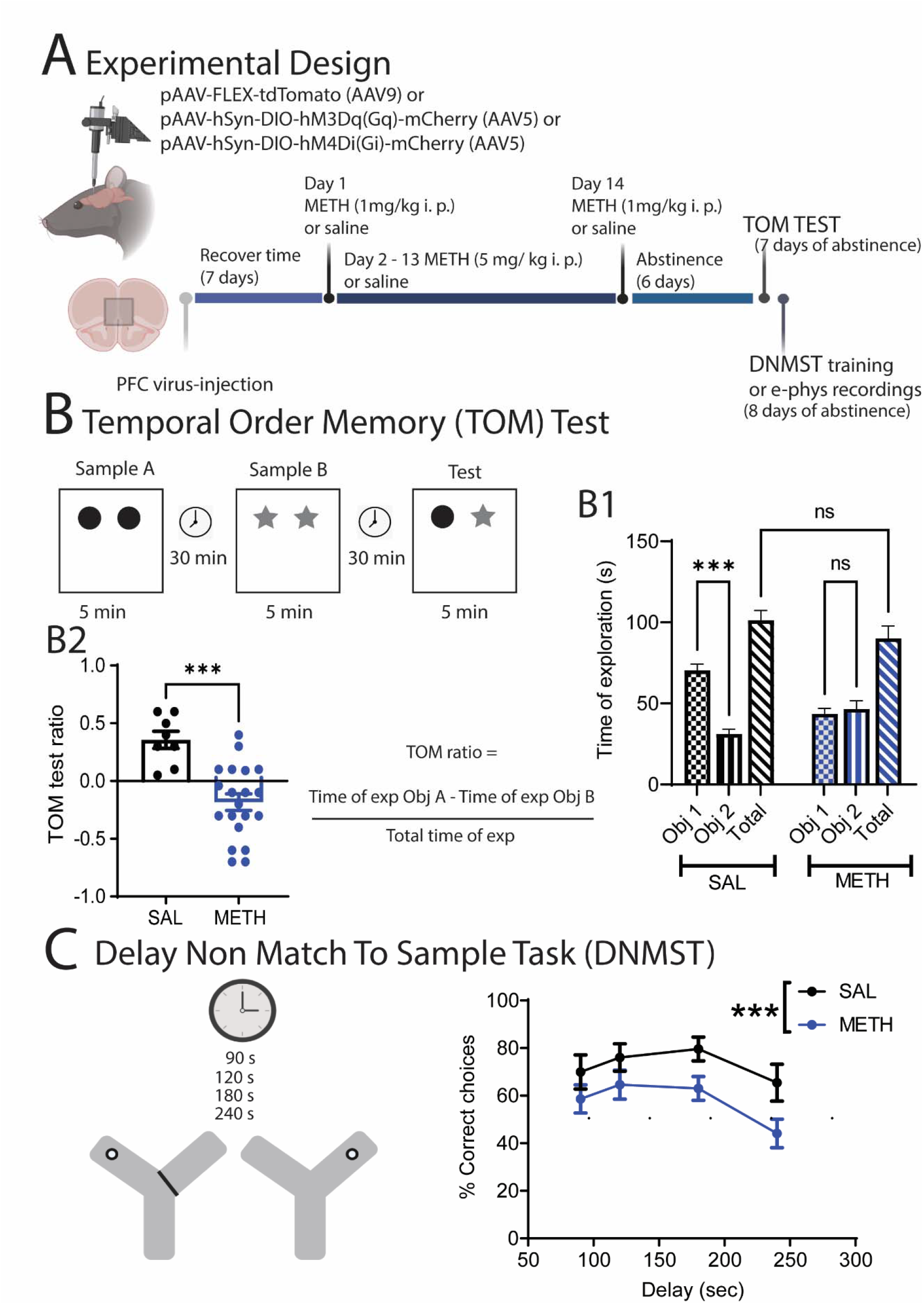
Experimental design and behavioral assessment. (A) Schematic illustrating the infusion of pAAV-FLEX-tdTomato, pAAV-hsyn-DIO-hM3Dq (Gq) or pAAV-hsyn-DIO-hM4Di (Gi) DREADDS in the PFC of PV Cre rats followed by repeated METH (or SAL) regime, cognitive tests and electrophysiological assessments. (B) Schematic representation of the Temporal Order Memory (TOM) test, (B1), Total time of exploration of the test and total time of exploration of each object, no significantly changes in total time of exploration of both groups [SAL Object A. 70.33 ± 3.8 vs SAL Object B. 31.00 ± 3.0; t-test *** p< 0.0001, METH Object A. 43.5 ± 3.4 vs METH Object B. 46.5 ± 5.1; t-test ns p = 0.6263, SAL total time 101.3 ± 6.0; METH total time 90.5± 7.6; t-test ns p = 0.3172] (B2) METH induces a reduction of the TOM test ratio [unpaired t-test **** p< 0.0001, Sal n=12, METH n=20] (C) Schematic representation of the Delay No Match to Sample Test, graph shows METH-induced impairment in working memory [Two-way ANOVA F (1,115) =11.3 *** p< 0.01, SAL n= 12; METH n=20]. The schematic illustration was created with BioRender.com.

### Intracranial surgeries

Adult male PV-Cre+ rats (270 – 320 g) were anesthetized with isoflurane. Viral vectors [Cre-dependent pAAV-Flex-tdTomato-AAV9; AAV-hSyn-DIO-HM3Dq-mCherry; AVV-hSyn-DIO-HM4Di-mCherry (1.5 μl – 3 μl)] were injected bilaterally into the mPFC using a stereotaxic apparatus. Injections were performed using a nanojector (Drummond Scientific Company, Broomall, PA, USA) and a glass pipette (1 μm tip diameter) using the following coordinates for prelimbic mPFC: PL-PFC [anterior-posterior (AP): +3.0- 3.2 mm; medial-lateral (ML): +/− 0.7 mm; dorsal-ventral (DV): −3.8 mm, Paxinos and Watson 1982]. Rats were single housed for 7 days following surgery and then moved to pair housing for METH or SAL administration.

### Cognitive assessments

#### Temporal Order Memory

(Fig. 1B). Animals were habituated for 2 consecutive days to the arena (45×35×40 cm) for 5 min in the absence of objects. During the first sample trial or phase 1, a single rat was placed into the arena and allowed to explore two identical objects for 5 min. Then the rat was placed into its home cage for 30 min. Phase 2 consisted of repeating the 5 min exploration but using a different pair of identical objects, before being placed again into its home cage for 30 min. Finally, during the test, the rat was placed again into the arena with one object from each sample (i.e., Object A and B), for 5 min (Fig. 1B). The criteria for evaluating behavioral exploration of the object was defined as: sniffing the objects and physically interacting with the object. The objects were counter balanced in all the groups evaluated. The discrimination ratio was calculated as the difference in time spent by each animal exploring object A compared with object B, divided by the total time spent exploring both objects during the five minutes of the test period. The total time of exploration was recorded and considered as a control for locomotor activity. In this test, the hypothetical ratio expected in a control animal is a positive ratio, meaning that the animal spent more time exploring the “oldest” object, since in order terms, the “oldest” object is novel compared with the most recent one (65). On the other hand, a ratio equal to or less than zero suggests a deficit in recognizing the most recent object, and means that animals spent equal time exploring the “oldest” and recent object or that they spent more time exploring the most recent object. All the phases of the TOM test (habituation, phase 1, phase 2 and test) were videotaped and the analysis and quantification were performed post hoc. All the objects used on the test were previously tested for preference by a control group of naïve rats. The rats didn’t show preference for any particular object.

In a group of PV-Cre rats (n=13), AAV-hSyn-DIO-HM3Dq-mCherry was injected bilaterally into the prelimbic PFC followed by SAL treatment. In a different group of PV-Cre rats (n=5) AVV-hSyn-DIO-HM4Di-mCherry was injected bilaterally into the prelimbic PFC, followed by repeated METH exposure. Clozapine N-oxide (CNO) was administered i.p, (10 mg/kg; (66)) and 30 minutes later the TOM test was performed. The TOM ratio was compared to animals that received vehicle solution.

#### Working Memory

(Fig. 1C). To test reference and WM, we used a modified version of a DNMST test (67). In brief, rats were food restricted (~90% of their original weight) and habituated to a Y maze for 3 days before the training phase. Training consisted of two sessions, an information run and a test run. In the information run, one arm was blocked forcing the animal into the open arm, where it received a sugar pellet. Subsequently, the blocked arm was open, and the rat was given a free choice of either arm. Rats received a sugar pellet for choosing the previously unvisited arm (correct choice), choosing the visited arm yield no reward. Left/right allocations for the information and test runs were pseudo-randomized over 10 trials/day with no more than 3 consecutive information runs to the same side. The inter-trial interval was 1 min. An entry to the arm was considered when the 4 paws of the rat and half length of the tail were on the selected arm. A correct choice was scored when the animal retrieved the sugar pellet. An incorrect choice was scored when the animal entered the previously visited arm. A no choice was scored when the animal did not enter the correct arm in a period of 2 minutes or entered a correct arm but did no retrieve the sugar pellet. After 3 d of training, ≥70 % correct choices for 2 consecutive days were required to proceed to the WM evaluation. For the WM test, rats were assigned randomly 10 trials with delays of 90, 120, 180 and 240 sec between the information run and the test run (the % of correct choices/delay was calculated for comparison).The performance in the delay component of the task was assessed for 2 consecutive days. All cognitive tests were performed during the light phase of the rats.

### Brain Slice Preparation

Animals were deeply anesthetized using isofluorane (Minrad Inc., Orchard Park, NY, USA), rapidly decapitated, and the brain extracted. Coronal slices of 300 μM thick containing mPFC were obtained with a vibratome (Leica VT 1200s, Leica Microsystems, Buffalo Grove, IL, USA) in ice-cod artificial cerebral spinal fluid [aCSF containing: 200 mM sucrose, 1.9 mM KCL, 1.2 mM Na2HPO4, 33 mM NaHCO3, 6mM MgCl2, 10 mM D-glucose, and 0.4 mM ascorbic acid (pH: 7.3 – 7.4, 305-310 mOsm)]. Slices were incubated in a hot bath at 31 °C for 1 h before recordings, with a physiological solution containing:120 mM NaCl, 2.5 mM KCl, 1.25 mM NaH2PO4, 25 mM NaHCO3, 4 mM MgCl2, 1 mM CaCl2, 10 mM D-glucose, and 0.4 mM ascorbic acid, oxygenated 95% o2/ 5% CO2 (pH: 7.3 – 7.4, 305-310 mOsm).

### Electrophysiological Recordings

Whole cell patch clamp was used to record neurons located in layers V - VI of the prelimbic mPFC identified using infrared-differential interference contrast optics and video microscopy. PV+FSIs were identified using a filter for tdTomato. Signals were recorded using an Axoclamp 2B (Molecular Devices), low-pass filtered at 3 kHz and digitalized at 10 kHz. Data acquisition was performed using Axograph-software (Axograph J. Clemens). For voltage-clamp recordings, electrodes (3-5 MΩ resistance *in situ*) were filled with a solution containing: 130 mM CsCl, 10 mM HEPES, 2 mM MgCl2, 0.5 mM EGTA, 2 mM Na2ATP, 0.3 mM Na-GTP, 2 mM QX-314, 10 mM phosphocreatine, (pH = 7.2 −7.3, 270- 280 mOsm). The membrane potential was clamped at −70 mV. Series resistance (Rs) was monitored throughout the experiment using a −5-mV pulse (50 ms duration) and recordings that exhibit changes in baseline > 30% were discarded. GABAergic-mediated events were pharmacologically isolated by adding CNQX (10 μM) and AP-V (50 μM) to the bath. Spontaneous Inhibitory Post-Synaptic Currents (sIPSC) recordings consisted on five sweeps/10 s. Evoked inhibitory synaptic currents (eIPSCs) and paired pulse (PP) recordings consisted in 8-10 repetitions at 0.5 Hz using the minimum amount of current needed to evoke a response. The sIPSCs were analyzed using MiniAnalysis software (Synaptosoft, Fort Lee, NJ USA). Synaptic currents were evoked using bipolar theta capillaries filled with saline solution placed in the mPFC around 300 μm from the patched neurons. Cells were recorded during baseline conditions using aCSF followed by 5-7 min of aCSF+ DA drugs. After 5-7 min of perfusion with the aCSF+DA drugs, the perfusion solution was switched to regular aCSF. The coefficient of variation (CV) was calculated as the standard deviation of the eIPSCS amplitude/amplitude of the eIPSCs (68). In order to normalize the CV data, we used 1/CV^2^.For current clamp recordings, a potassium –based internal solution was used. Parvalbumin-positive fast spiking interneurons (PV+FSI) were identified by tdTomato fluorescence and electrophysiological signature(29). tdTomato fluorescent neurons that did not display the electrophysiological characteristics of fast spiking interneurons were excluded from the analysis. Current steps of 20 pA from −100 to 300 pA (1 sec duration) were injected to generate I/F and input resistances curves.

Excitatory/inhibitory ratio (E/I) was recorded using a potassium-based internal solution (in mM): 120 K-gluconate,10 NaCl, 10 KCl, 0.5 EGTA, 2 Na2ATP, 0.3 Na-GTP, 10 HEPES and 10 phosphocreatine, pH 7.2-7.35 and 270-280 mOsm.. Evoked PSCs at −70 mV and 0 mV were recorded and the ratio calculated. Control experiments recording a 0 mV in SAL rats (n=3) showed that application of picrotoxin at this membrane potential blocked all spontaneous activity.

### Immunohistochemistry

Rats were given an overdose of ketamine/xylazine (150mg/kg and 2.9mg/kg respectively) diluted in 0.9% saline, then perfused transcardially with 4% paraformaldehyde (PFA). The brains were post-fixed overnight in 4% PFA, cryoprotected in 30% sucrose for 3 days and sectioned at 40um using a cryotome. Sections were washed in PBS 1X, incubated in blocking solution (5% normal donkey serum, 1% Bovine Serum Albumin, 0.2% glycine, 0.2% lysine, 0.3% Triton X-100 diluted in PBS 1X) under shaking for 1 hour at room temperature, incubated with primary antibodies diluted in blocking solution (mouse anti-parvalbumin (1:800) from Millipore #MAB1572; rabbit anti-DsRed (1:800) from LivingColors #632496) overnight at 4C under shaking, washed 3 times 10 minutes in PBS 1X under shaking at room temperature, incubated with secondary antibodies diluted in blocking solution (AlexaFluor 488 donkey anti-mouse (1:500) from Fisher #A21202; AlexaFluor 594 donkey anti-rabbit (1:500) from Fisher #A21207) for 1 hour and 30 minutes at room temperature under shaking and protected from light, washed 3 times 10 minutes in PBS 1X under shaking and finally mounted in ProLong Gold Antifade Mountant (Invitrogen #P36931). Sections were imaged using a Zeiss confocal: z-stacks were made at 20X in the mPFC. All quantifications were performed using ImageJ.

| Reagent or resource | Source | Identifier |
| --- | --- | --- |
| ***Antibodies*** |  |  |
| Rabbit anti-PV | Abcam | ab11427/ GR3317380-1 |
| Goat Anti-Rabbit IgG h&L (Alexa Fluor 488) | Abcam | Ab150077/ GR3279569-1 |
| ***Bacterial and Virus Strains*** |  |  |
| pAAV-FLEX-tdTomato (AAV9) | Addgene | Addgene.org/28306 |
| pAAV-hSyn-DIO-hM3Dq(Gq)-mCherry (AAV5) | Addgene | addgene.org/44361 |
| pAAV-hSyn-DIO-hM4Di(Gi)-mCherry (AAV5) | Addgene | addgene.org/44362 |
| ***Software*** |  |  |
| Axograph |  |  |
| Prism GraphPad |  |  |
| Image J | NIH | https://imagej.nih.gov/ij/ |
| Mini analysis |  |  |
| ***Drugs*** |  |  |
| Methamphetamine | Sigma | CAS 51-57-0 |
| Clozapine N-oxide (CNO) dihydrochloride (water soluble) | Hellobio | HB6149 |
| SKF 81297 | Sigma | CAS 253446-15-0 |
| SCH 22390 | Sigma | CAS 125941-87-9 |

### Data Analysis and Statistics

Electrophysiological data analysis was performed using Axograph software® (J. Clemens) and MiniAnalysis® (Synaptosoft, Fort Lee, NJ USA). Electrophysiological and behavioral data were analyzed using GraphPad software. Data are presented as mean ± SEM. Normality was determined using Shapiro-Wilk test: for normal data, we used Student’s t test for comparison of two independent groups. In the case of multiple comparison, we used ANOVA followed by Tukey’s post-hoc test. For comparison of intrinsic excitability, a Two-way ANOVA was used.

## Results

### Chronic METH administration produces cognitive deficits

To examine the effect of chronic, daily METH administration and abstinence on working memory behavior in rats, we injected rats with METH or Saline (SAL) for 14 consecutive days (i.p., see methods and Fig 1A) followed by 7-10 days of home-cage abstinence. We then tested the rats in two PFC-dependent cognitive behavior tests, the Temporal Order Memory test (TOM) and the Delayed Non-Match to Sample Task (DNMST)(65) (Fig. 1B,C). In the TOM test, rats were allowed to interact with a pair of identical objects (Object A). After a 30-minute break in the home cage, the rats were allowed to interact with a different pair of identical objects (Object B). After a second 30-minute break, they were allowed to interact with Object A and Object B together. As expected, the saline-treated rats showed a preference to interact with the non-recent object A (Fig. 1 B1, SAL Object A. 70.33 ± 3.8 vs SAL Object B. 31.00 ± 3.0; t-test p< 0.0001). In contrast, METH-treated rats spent same time exploring both objects (Fig. 1 B1, METH Object A. 43.5 ± 3.4 vs METH Object B. 46.5 ± 5.1; t-test ns p = 0.6263), and there was a significant difference by METH treatment (Fig. 1 B2, unpaired t-test p< 0.0001, Sal n=12, METH n=20), indicating a likely deficit in short-term spatial memory. There was no significant effect of METH treatment on total time interacting with the objects (Fig. 1 B1 SAL total time 101.3 ± 6.0; METH total time 90.5± 7.6; t-test ns p = 0.3172), so METH didn’t appear to alter exploratory drive or motor ability. In the DNMST, a working memory task where rats must remember the most recent reward location to make the subsequent non-matching location choice, METH-treated rats showed a significant decrease in the percentage of correct choices [Two-way ANOVA F (1,115) =11.3 p< 0.01, SAL n= 12; METH n=20; Fig 2C], compared to SAL controls. Together, these data demonstrate that chronic METH exposure in rats produces significant working memory deficits, similar to the cognitive symptoms in individuals suffering from METH use disorder.

**Figure 2.**
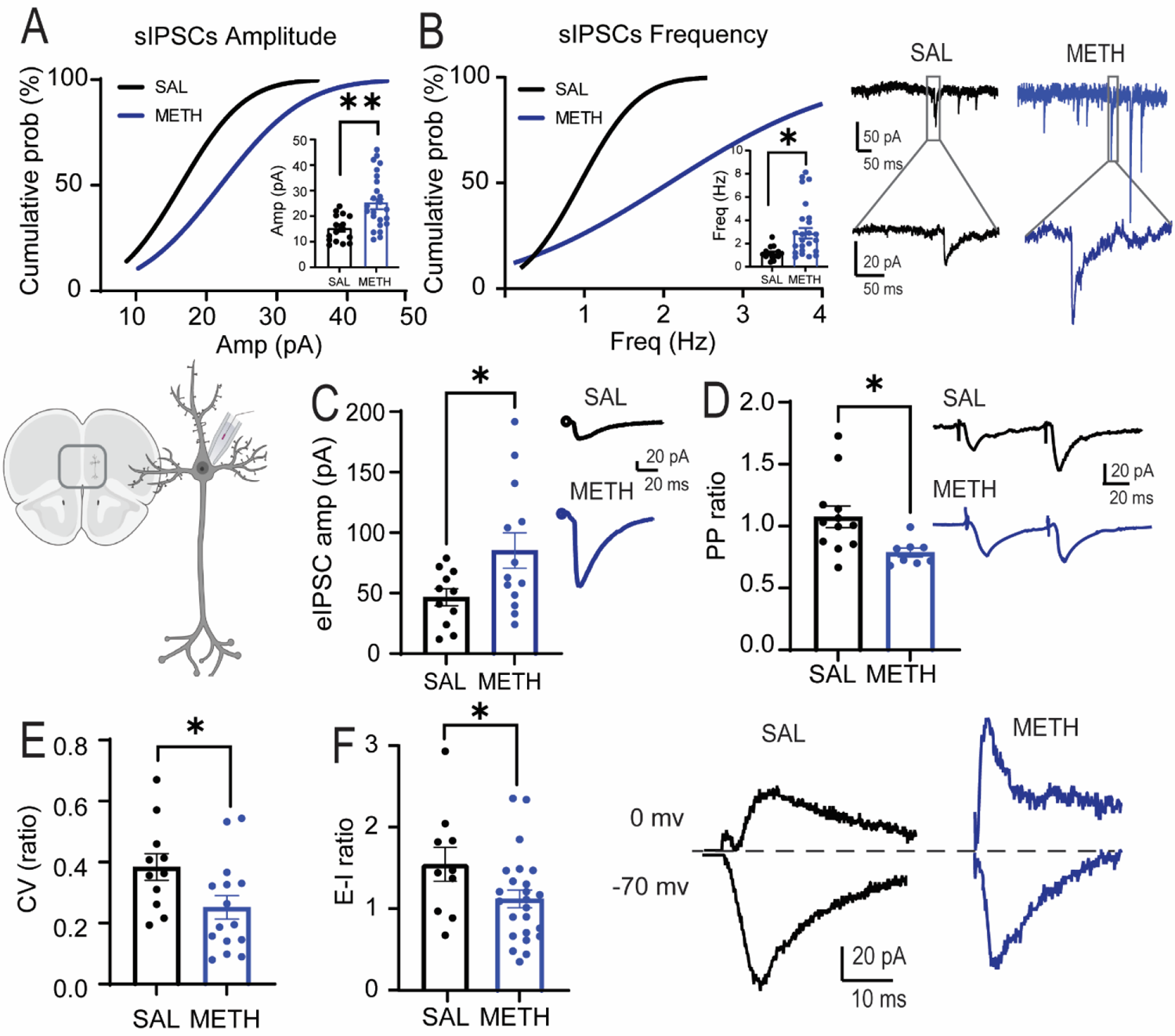
Repeated METH increases GABA transmission. (A) Repeated METH evokes a significant increase in the amplitude of sIPSCs recorded in PNs [SAL = 1.18 ± 0.13 Hz, n= 18/8; METH = 3.1 ± 0.2 Hz, n=25/8, Student’s t test, ** p = 0.0028]. (B) Repeated METH elicits a significant increase in the frequency of sIPSCs [SAL = 14.35 ± 1.3 pA n=18/8; METH = 24.4 ± 2.1 pA n=25/8, Student’s t test * p = 0.0124]. Representative traces are shown to the right. (C) Repeated METH elicits a significant increase in amplitude of eIPSCs [SAL = 46.56 ± 6.9 pA n= 11/7; METH = 85.22 ± 14.6 pA n=15/9; Student’s t test * p = 0.0353]. Representative illustration of recordings is shown to the left and representative traces to the right. (D) Repeated METH elicits a significant paired pulse depression [SAL= 1.07 ± 0.08, n=12/6; METH= 0.78 ± 0.04, n=8/6; Student’s test, * p = 0.0188] (E) Repeated METH elicits a significant decrease in eIPSCs coefficient of variation [SAL= 0.38 ± 0.04, n=11/7; METH= 0.25 ± 0.03, n=15/9; Student’s test, * p = 0.0337]. (F) Repeated METH elicits a significant decrease in E-I ratio measured in PNs [SAL = 1.5 ± 0.2 n=10/7; METH = 1.0 ± 0.1 n= 25/10; Student’s test, * p = 0.042]. Representative traces are shown to the right. All results are reported as mean (X) ± standard error (SE). N=number cells/rat. The schematic illustration was created with BioRender.com.

### Repeated METH treatment increases inhibitory transmission in PFC deep layers

Working memory tasks, like TOM and DNMST, are dependent on PFC function (65,67,69). To test whether repeated, daily METH administration in male adult rats produced electrophysiology changes in PFC synaptic transmission after abstinence that are similar to a hypofrontal state, we performed patch-clamp recordings of deep-layer (layer V-VI) pyramidal neurons (PNs) in acute slice preparations. Repeated METH exposure significantly increased both the amplitude (Fig. 2A, unpaired student’s t-test, p=0.002) and frequency (Fig. 2B; unpaired student’s t-test p = 0.01) of spontaneous inhibitory postsynaptic currents (sIPSCs) on PNs, indicating that chronic METH exposure increased PFC deep-layer GABAergic synaptic transmission. Consistent with the sIPSC findings, the METH-treated rats also showed a significant increase in the amplitude of evoked IPSCs (eIPSCs) (Fig. 2C; unpaired student’s t-test p = 0.03). Using paired-pulse ratio analysis (PPR, 50 ms interstimulus interval), we found that METH-treated animals showed a decrease in PPR (Fig. 2D, unpaired student’s t-test, p = 0.01), suggesting an increase in GABA release properties of inhibitory inputs onto PNs. Chronic METH-treated rats also displayed a significantly reduced eIPSC coefficient of variation (CV) (Fig. 2E, unpaired student’s t-test, p = 0.03), consistent with an increase in GABAergic presynaptic probability of release (P_r_) and/or the number of synaptic contacts (68). Finally, we assessed whether the increased inhibition onto PNs altered the E-I ratio by recording both eEPSCs (−70mV) and eIPSCs (0mV) from the same PNs, and we observed a significant decrease in the E-I ratio (Fig 2F, unpaired student’s t-test, p = 0.04). Taken together, our data indicate that chronic METH administration plus one-week of homecage abstinence increases GABAergic synaptic transmission onto prelimbic PNs and alters E-I ratio, consistent with hypofrontal circuit function.

### Dopamine D1 receptors are required for METH-induced changes in PRL inhibitory synaptic transmission

METH produces a strong increase in dopamine release(46,47,70) and dopamine receptor signaling (71), and D1 dopamine receptors (D1R) are expressed in PNs and PV+FSIs interneurons in the PFC(56,72,73). To test whether D1R signaling influences the chronic METH-induced increase in GABAergic synaptic transmission onto PFC PNs, we prepared PFC acute slices from saline- or METH-treated rats and assessed the effects of the selective D1R antagonist SCH 23390 (10 μM). While the D1R antagonist had no significant effects on sIPSC, eIPSCs, or PPR of PNs from saline-control treated rats (Supplemental Fig. 1 B1-4), it significantly decreased the amplitude (paired student’s t-test p = 0.0003, Fig 3A) and frequency (paired student’s t-test p= 0.03; Fig. 3B) of sIPSCs, and the eIPSC amplitude (Fig.3D, paired student’s t-test, p = 0.02) in slices from METH-treated rats. Interestingly, SCH 23390 did not alter the METH-induced changes in PPR (Fig.3E, paired student’s t-test, ns p = 0.6) which suggests that the METH-induced increases in s/eIPSCs are largely independent of the changes in presynaptic short-term plasticity. Finally, to test whether activation of D1Rs is sufficient to enhance IPSCs onto deep-layer PNs, we also treated slices from saline-treated rats with the D1R agonist SKF 81297 (5 μM). However, acute pharmacological activation of D1R signaling did not alter sIPSCs, eIPSCs, or PPR (Supp Fig 1 A1-4), suggesting that it is necessary, but not sufficient, for METH-induced changes in GABAergic synaptic transmission.

**Figure 3.**
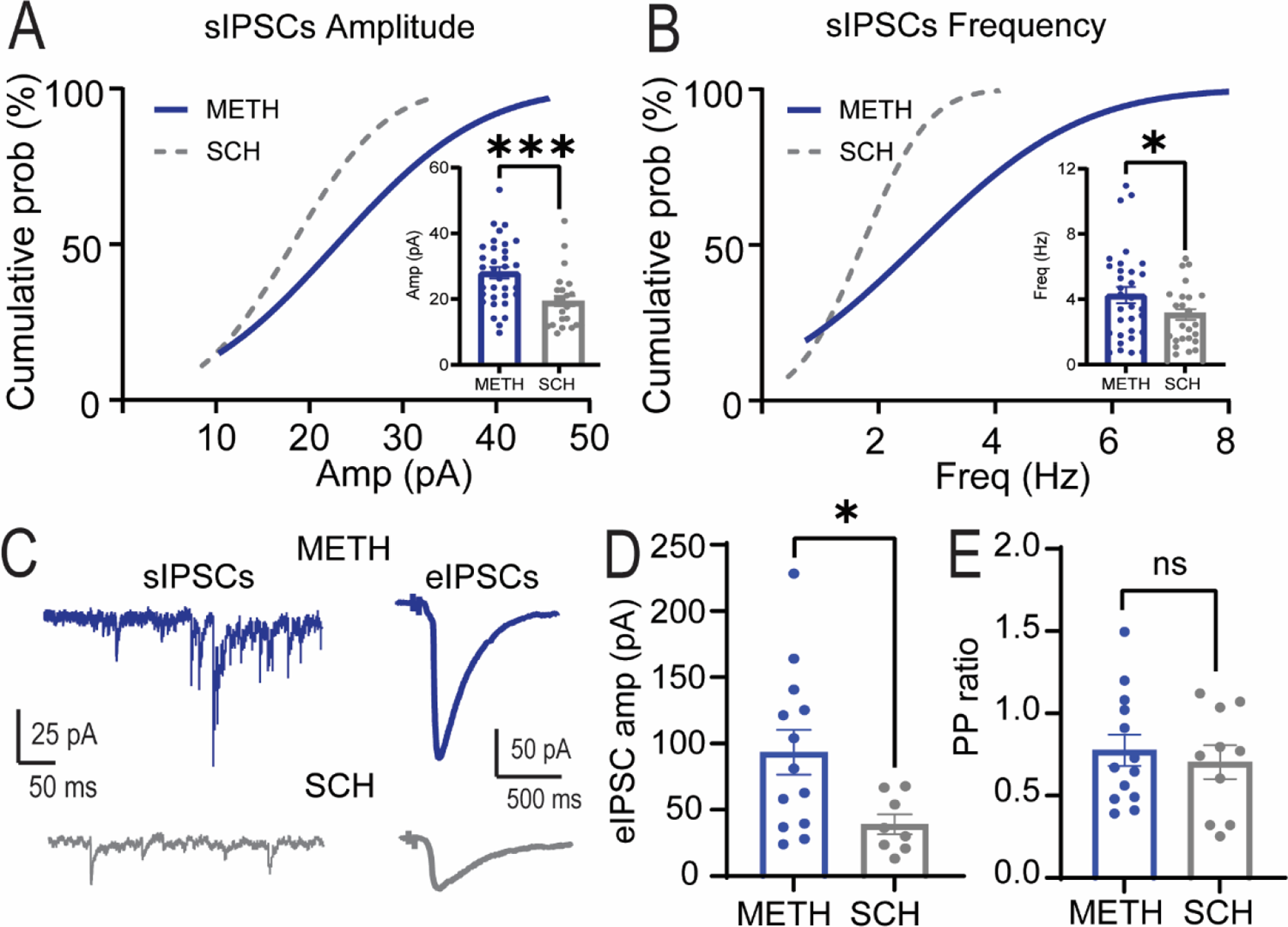
Bath application of the D1 antagonist SCH 23390 decreases repeated METH– mediated increases in inhibitory activity in PNs recorded in mPFC. (A) Bath application of the selective D1R antagonist SCH 23390 (10 μM) produces significant decreases in the sIPSCs amplitude of rats treated with repeated METH [METH = 24.3 ± 2.5 pA; SCH = 19.0 ± 1.6 pA paired Student’s t test *** p = 0.0003; n=18/7]. (B) SCH 23390 produces significant decreases in the frequency of sIPSCs recorded from rats treated with repeated METH [METH = 3.3 ± 0.5 Hz; SCH = 1.9 ± 0.2 Hz; paired Student’s t test, * p= 0.0301, n = 18/7]. (C) Representative traces of sIPSC and eIPSC recorded in PNs of rats treated with repeated METH before and after bath perfusion of SCH 23390 (D) SCH 23390 bath perfusion elicits a significant decrease in the amplitude of the eIPSCs [METH= 90.2 ± 20.7 pA; SCH = 46.8 ± 10.5 pA, paired Student’s t test, * p = 0.0266; n=13/6]. (E) Bath administration of SCH 23390 did not alter the PP of PNs recorded in rats treated with repeated METH [paired student’s t-test, ns p = 0.6106 n= 13/6].

### Chronic METH administration alters the intrinsic excitability of PV-FSIs

Since chronic METH administration increased inhibitory synaptic transmission onto PFC PNs, we tested whether chronic METH altered PV+FSIs function, a major contributor to PFC inhibitory transmission. To this end, we injected Cre-dependent tdTomato-expressing virus into the PFC of PV-Cre rats (Fig. 4A), and after 3 weeks, we found that 93% of the tdTomato-positive cells were parvalbumin-positive Fig. 4B). Using whole-cell current clamp configuration, we recorded PV-FSIs from chronic saline or METH-treated animals. Compared to saline-treated rats, chronic METH treatment produced a significant increase in the frequency of action potentials elicited by a range of injected currents [Fig. 4C, Two-way repeated-measures ANOVA, F (1,162) =56.6, p< 0.0001], indicating that METH produces an increase in PFC PV-FSI intrinsic excitability. Of note, there were no significant differences in the input resistance of PV+FSIs between SAL and METH groups (SAL: 446.2 ± 98.1 MΩ; METH: 311. 4± 76.3 MΩ), suggesting that the increase in PV+FSIs excitability is not mediated by METH-induced alterations in PV+FSIs constitutive channels.

**Figure 4.**
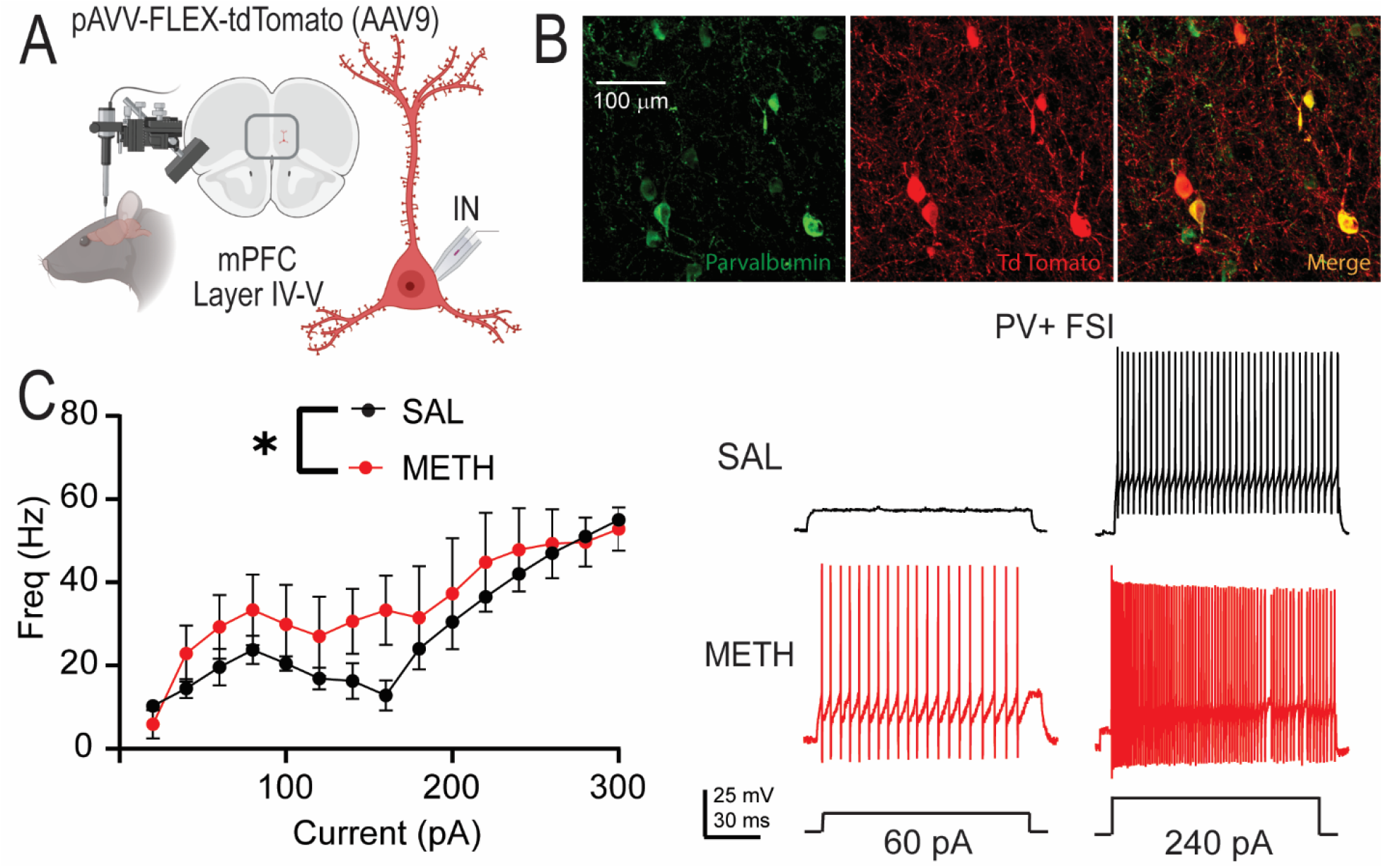
Repeated METH alters the firing frequency of interneurons in the mPFC. (A) Schematic representation of intracranial injection of pAVV-FLEX-tdTomato in the mPFC to label parvalbumin-positive interneurons (PV+FSIs). (B) Colocalization of PV (green) and pAVV-FLEX-tdTomato (red) show selectivity for labeling PV+FSIs (yellow). (C) Rats treated with repeated METH exhibit a significant increase in PV+FSIs intrinsic excitability [Two-way ANOVA, F (1,162) =56.6, **** p < 0.0001, SAL n= 5/4; METH n= 7/5]. Representative traces are shown to the right. The schematic illustration was created with BioRender.com.

### Chemogenetic reduction of PFC PV+FSI activity rescues METH-induced cognitive deficits

To test whether an increase in PV+FSI function is sufficient to mimic METH-induced deficits in working memory, we injected the PFC of saline-treated PV-Cre rats with a Cre-dependent excitatory DREADD (AAV5-DIO-GqDREADD). After 3 weeks of virus expression, we observed strong co-localization of GqDREADD/mCherry with endogenous parvalbumin (Fig. 5A right panel), confirming the selective expression of DIO-GqDREADDs in PV+FSIs. The GqDREADD-expressing rats were injected with either vehicle or CNO (10 mg/kg; i.p.) to activate PFC PV+FSIs 30 minutes before testing the temporal order memory task. Compared to animals that received vehicle, the CNO-treated rats showed a significant decrease in the TOM ratio (Fig 5A, One-way ANOVA F(3,41)=10.21, Tukey’s *post hoc* test, p< 0.0001), indicating that increasing PFC PV+FSI activity is sufficient to reduce working memory function. To determine whether the increased PFC PV-FSI activity is necessary for the chronic METH-induced WM deficits, we injected chronic METH-treated PV-Cre rats expressing a Cre-dependent inhibitory DREADD virus (AAV2-DIO-GiDREADD) in the prelimbic cortex. We confirmed that 87% of the GiDREADD/tdTomato-expressing cells colocalized with PFC parvalbumin (Fig. 5B right panel). As expected, vehicle control-treated rats after chronic METH administration displayed significant deficits in the TOM ratio (Fig. 5B); whereas, CNO-treated rats after chronic METH displayed a TOM ratio similar to the saline-treated rats and significantly different than the METH + vehicle group (Fig 5B, one-way ANOVA F(2,27)=15.93, *Tukey’s post hoc*, p< 0.0001]. Together, these findings suggest that increasing PFC PV+FSIs activity is both necessary and sufficient for chronic METH-induced deficits in WM performance.

**Figure 5.**
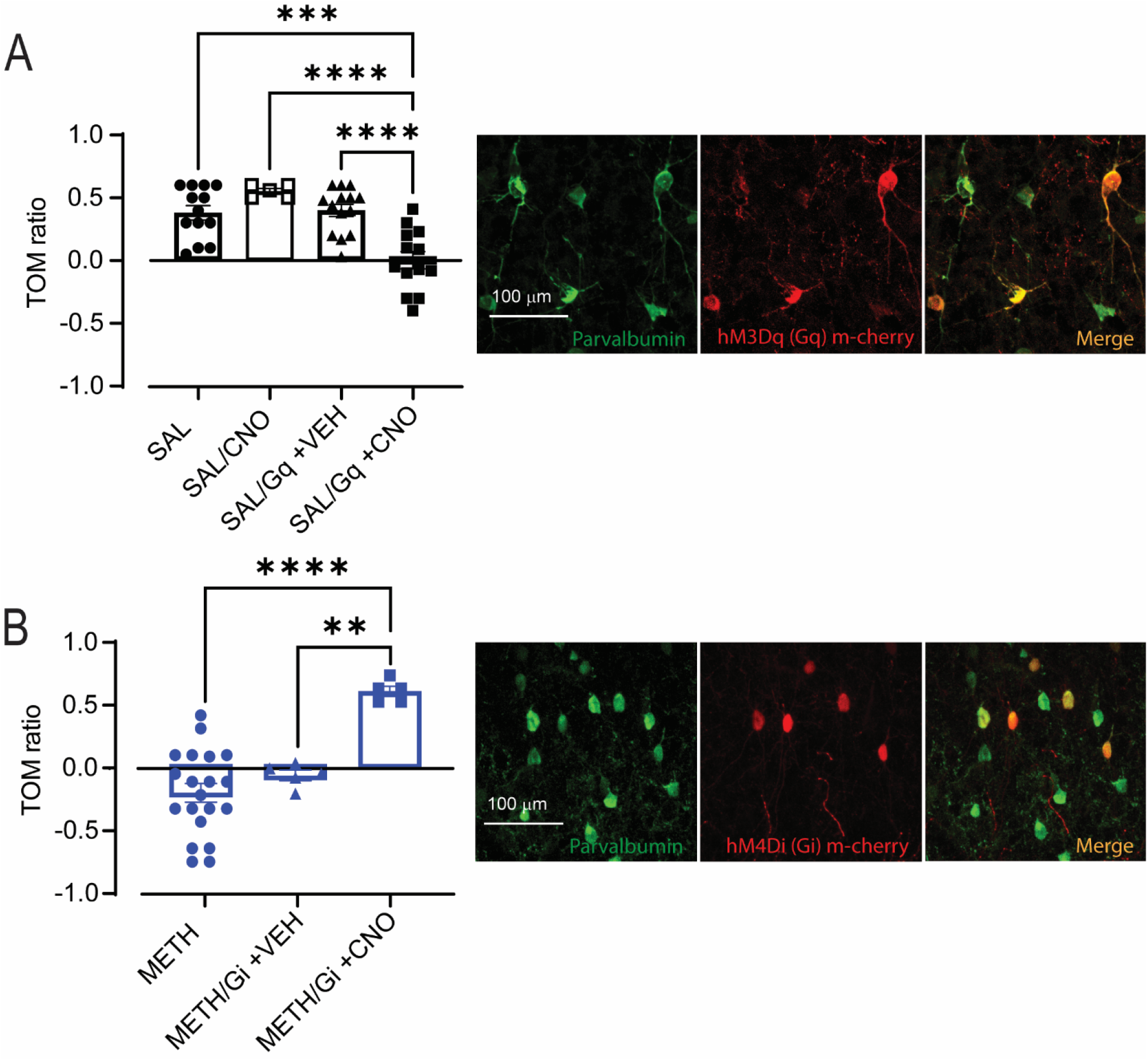
Chemogenetic manipulation of PV+FSIs. (A) Temporal Order Memory Test (TOM) with the infusion of DREADDs in the mPFC. Activation of PV+FSIs in SAL rats with Gq-DREADD replicates the deficits in TOM elicited by repeated METH [One-way ANOVA F(3,41)=10.21, Tukey’s *post hoc* test, *** p< 0.001, **** p< 0.0001]. Colocalization of PV and Gq-DREADD (right panel). (B) On the other hand, inactivation of PV+FSIs with Gi-DREADD in rats treated with repeated METH ameliorates the deficits in TOM elicited by repeated psychostimulant administration [One-way ANOVA F(2,27)=15.93, *Tukey’s post hoc*, **p< 0.01, **** p< 0.0001]. Colocalization of PV and Gi-DREADD (right panel). Temporal Order Memory ratio was calculated as the difference between object number 1 – object number 2, divided into the total time of exploration.

## Discussion

We found here that chronic METH administration followed by 7-10 days of abstinence elicited significant reductions in WM function and long-lasting increases in PFC GABAergic transmission. D1R signaling was required for the METH-induced increase in PFC GABAergic synaptic transmission. In addition, chronic METH increased PFC PV+FSI excitability, and chemogenetic reduction of PFC PV+FSIs activity rescued the METH-induced WM deficits, suggesting that a maladaptive increase in PV+FSIs activity, and possibly other PFC interneurons, underlies the METH-induced hypofrontality and WM dysfunction.

In our study, we noted multiple changes in GABAergic synaptic transmission and cell function following repeated METH administration, including: 1) an increase in the amplitude and frequency of sIPSCs onto deep-layer PNs in the PFC, 2) an increase in the amplitude of eIPSCs, 3) a decrease in E-I ratio, 4) the presence of inhibitory paired-pulse depression, 5) a decrease in eIPSCs CV, and 6) an increase in PV+FSI intrinsic excitability. The paired-pulse depression and decrease in eIPSC CV are consistent with an increase in GABAergic presynaptic function (68,74), although a change in CV could also reflect a change in GABAergic synapse number, which could also contribute to the observed changes in IPSCs. Future studies will be important to assess that possibility, and to investigate the relative contributions of various GABAergic interneurons to the noted changes. Nonetheless, our studies suggest that enhanced PV+FSIs activity and excitability could underlie, at least in part, the METH-induced electrophysiology and behavioral effects.

Studies have shown that in layer V of the PFC, only PV neurons show increases in FOS expression following an amphetamine regime (75) and chronic cocaine elicits an increase in PV+FSIs excitability(76). Similarly, Shi and colleagues(77) showed that in mice, repeated cocaine administration increases mIPSCs and our previous results showed that a repeated cocaine regime increases the D1R-mediated amplitude of eIPSCs(78). However, to the best of our knowledge, this is the first report describing the effects of repeated METH on the activity of PV+FSIs in PFC.

Our results show that the selective D1R antagonist SCH 23390 significantly decreased METH-enhanced amplitude and frequency of sIPSCs in chronic METH-treated rats, but without affecting synaptic measures in SAL-treated rats. Furthermore, the D1R antagonist significantly decreased the amplitude of eIPSCs and induced a trend for an increase in E-I ratio, suggesting that blockade of D1R decreased GABAergic synaptic transmission. However, increases in amplitude of eIPSCs could also suggest an increase in GABAR and/or changes in GABAR affinity, and the lack of effects of the D1 antagonist in paired-pulse ratio might suggest that the METH-induced increases in eIPSCs and sIPSCs are not caused by changes in presynaptic probability of release, but rather by increases in GABAergic cell firing, PN GABAergic synapse strength, and/or the number or sub-cellular location of GABAergic synapses on PNs. Also, the effects of the D1R antagonist on the inhibitory synaptic activity in METH-treated rats, but not SAL-treated rats, might suggest that chronic drug exposure produces D1R signaling sensitization. Future studies investigating D1 signaling in PNs and PV+FSIs will be important to test this possibility. However, it is worth noting that increased D1R signaling is not sufficient to explain our observations of increased GABAergic function since treatment of SAL-treated rat slices with the selective D1R agonist, SKF 81297, did not mimic the effects of repeated METH administration, suggesting that other cellular events are needed to produce the changes in inhibitory synaptic transmission in METH-treated animals.

Cognitive deficits produced by METH have been extensively reported,(4,5,11,79–82), and our results confirm that repeated METH elicits temporal order memory and working memory deficits. More importantly, our results are the first to demonstrate that activation of Gq-linked receptors in PFC PV+FSIs of SAL treated rats mimics the cognitive deficits produces by repeated METH, while activation of Gi-linked DREADDs in PFC PV+FSIs of repeated METH rats ameliorates the cognitive deficits. These results suggest that maladaptive changes in PFC PV+FSI activity is necessary and sufficient for METH-induced cognitive deficits, likely by shifting the PFC E-I balance. Moreover, gamma oscillations are key components of cognitive processes (83,84) and involve the reciprocal interaction between PV+FSIs and principal pyramidal cells (85). The predominant mechanism underlying this type of rhythm is the phasic excitation of interneurons following spike generation in PNs(86). Gamma oscillations are thought to help to organize ensembles of synchronously firing neurons in various brain regions, and may play a key role in top-down mechanisms controlling cortical processing, such as attention(87), working memory(88) and decision-making (89). Augmented firing of PV+FSIs could affect gamma oscillations and perturb normal information processing in PFC and associated cognitive processes.

In summary, our results show that repeated METH administration alters PV+FSI excitability, GABAergic synaptic transmission in the mPFC, and E-I ratio onto deep-layer mPFC PNs. Moreover, we show that D1R signaling is required for some of the METH-induced changes in PFC GABAergic synaptic transmission, and that modulation of PV+FSI activity is necessary and sufficient for METH-induced deficits in PFC-dependent working memory.

## Acknowledgments

This work was supported by funding from NIDA R21 DA045889 and R01 DA054589 (A.L.), R01 DA032708 and P50 DA046373 (C.W.C.), and NIDA/ORWH U54 DA016511 in partnership with MUSC (M.A.R.).

The authors report no biomedical financial interests or potential conflicts of interest.

