## Supplementary figure 1 for "Chronic methamphetamine administration produces cognitive deficits through augmentation of GABAergic synaptic transmission in the prefrontal cortex"

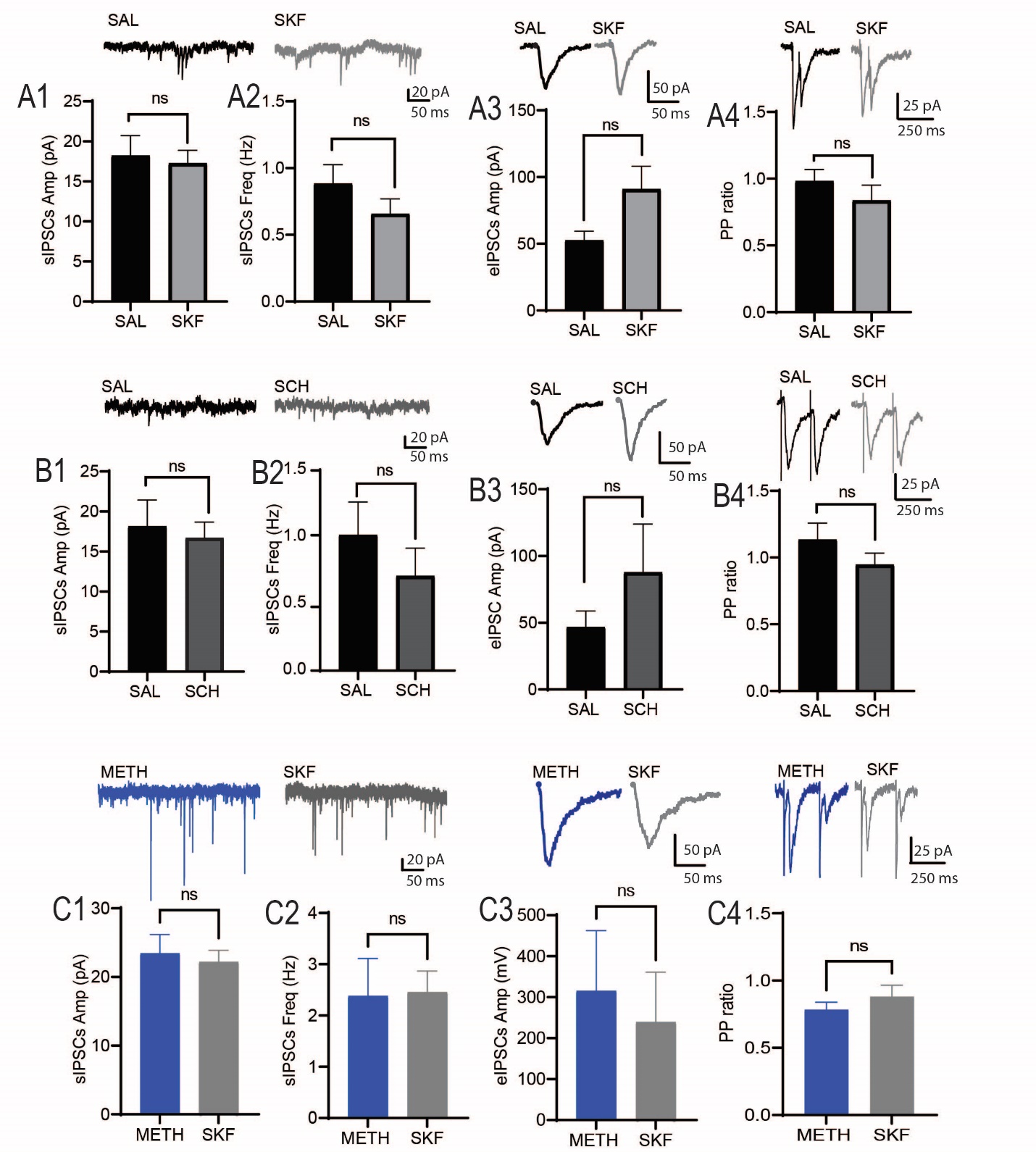
Figure S1. **Effects of** **Bath application of the D1 antagonist SCH 23390 (10µM) or D1 agonist SKF 81297 (5 µM) in Saline treated rats and Effects of Bath application of the D1 agonist SKF 81297 (5 µM) in METH treated rats** (A1) Bath application of the selective D1 agonist SKF 81297 did no alter the amplitude of sIPSCs in SAL treated rats (SAL= 18.2 ± 2.4 pA; SKF 81297 = 17.2 ± 1.6 pA, n= 12/6). (A2) Bath application of SKF 82397 do not later the frequency of sIPSCs in SAL treated rats (0.88 ± 0.14 Hz; SKF 81297= 0.65 ± 0.11 Hz, n=12/6). (A3) Bath administration of SKF 81297 elicits a trend for increase in the amplitude of eIPSCs in SAL treated rats (SAL = 52.6 ± 6.8 pA; SKF 81297= 90.9 ± 17.1 pA, paired t- test p= 0.06, n= 11/5). (A4) Bath administration of SKF 81297 did not alter PP ratio in SAL treated rats (SAL = 0.9 ± 0.08; SKF = 0.8 ± 0.11, n=10/5). (B1) Bath application of the selective D1 antagonist SCH 23390 to brain slices obtained from SAL rats did not alter amplitude of sIPSCs (SAL = 18.1 ± 3.2 pA; SCH = 16.7 ± 1.9 pA, n=9/6). (B2) Bath application of SCH 23390 to brain slices obtained from SAL show a trend for decreasing frequency of sIPSCs (SAL = 3.3 ± 0.8 Hz; SCH = 2.3 ± 0.6 Hz n=10/6, paired t-test p= 0.08). (B3)	Bath application of SCH 23390 did not alter the amplitude of eIPSCs in SAL rats (SAL =46.8 ± 11.9 pA; SCH = 87.8 ± 36 pA, n=8/6). (B4) SCH 23390 elicited a trend for PP	depression in PNs of SAL rats (SAL =1.1 ± 0.1; SCH = 0.9 ± 0.08, n=9/6, paired t-test p= 0.07). (C1) Bath application of the selective D1 agonist SKF 81297 did no alter the amplitude of sIPSCs in METH treated rats (METH= 23.43 ± 2.7 pA; SKF 81297 = 22.2 ± 1.6 pA, p = 0.722 n= 6/3). (C2) Bath application of SKF 82397 do not later the frequency of sIPSCs in METH treated rats (METH = 2.3 ± 0.7 Hz; SKF 81297= 2.4 ± 0.4 Hz, n=6/3). (C3) Bath administration of SKF 81297 elicits a trend for increase in the amplitude of eIPSCs in METH treated rats (METH = 315.1 ± 147.1 pA; SKF 81297= 238.9 ± 122.3 pA, paired t- test p= 0.7, n= 5/3). (C4) Bath administration of SKF 81297 did not alter PP ratio in METH treated rats (METH = 0.78 ± 0.05; SKF = 0.87 ± 0.08, n=5/3).
